# Methanotrophic acetogenesis drives a novel pathway of arsenic mobilization in reducing groundwaters

**DOI:** 10.64898/2025.12.23.696296

**Authors:** NM Bassil, W Xiu, LA Richards, BE van Dongen, DA Polya, HM Guo, AJ Probst, JR Lloyd

## Abstract

Geogenic arsenic (As) contamination in groundwater, widely used as drinking water, poses a global health risk, yet the microbial pathways linking electron donor oxidation to the reduction of As-bearing Fe(III) oxyhydroxides remain poorly understood. Here, we deployed Fe(III) (oxyhydr)oxide-coated pumice stones in high-As, high-methane groundwater in Cambodia for 270 days to capture planktonic metal-reducing microbial communities *in-situ*. These were used to inoculate anaerobic microcosms with methane or volatile fatty acids (VFAs) as electron donors over 200 days. Genome-resolved metagenomics revealed that methane oxidation via reverse methanogenesis led to acetate production, which in turn provided the primary electrons for Fe(III) and As(V) reduction in the microcosms, resulting in As(III) release. Our findings highlight an indirect coupling between methane oxidation and arsenic mobilization, with acetate as the key intermediate. This study offers new insights into the role of methane in subsurface biogeochemical cycling and its implications for arsenic contamination in groundwater systems.

**Synopsis:** The study uses a novel in situ sampling procedure, coupled with metagenomic analysis, to identify anaerobic methane oxidation producing acetate, as a potentially important pathway to generate electron donors for microbial Fe(III) and As(V) reduction in aquifer sediments.

**Graphical abstract:** 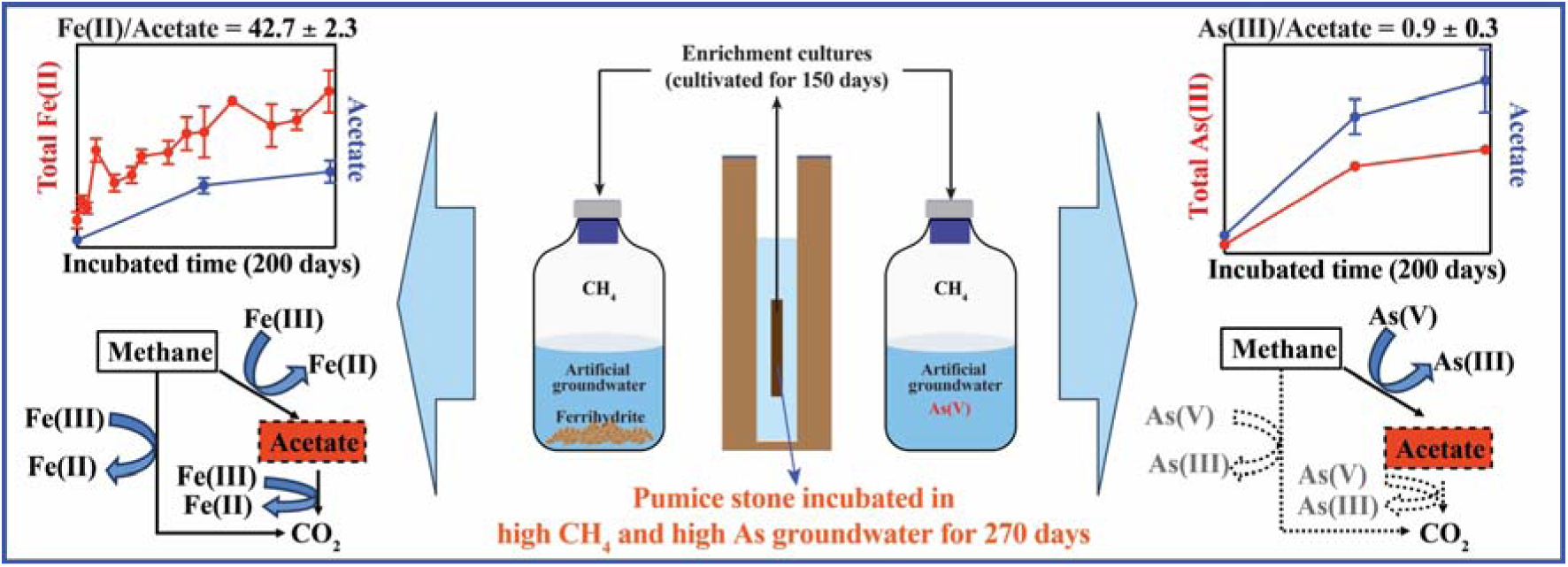

## Introduction

Geogenic high arsenic As groundwater (provisional World Health Organization guideline value for drinking water 10 μg/L) is a global issue ^1,2^, particularly threatening many millions of people in South and Southeast Asia ^3^ through increased risks of cancers ^4^ and cardiovascular diseases ^5^ amongst many other detrimental health outcomes ^6^. Arsenic mobilization from aquifer sediments is closely linked to the biogeochemical cycling of C, N, S, and Fe ^7–11^. Key mechanisms include the microbial reductive dissolution of As-bearing Fe(III) (oxyhydr)oxides ^9^, and direct As(V) reduction to the often more mobile As(III) via either detoxification (*ars* operon) or respiratory (*arr* operon) pathways ^12,13^.

Microbial Fe(III) and As(V) reduction can be fueled by diverse electron donors, including dissolved organic matter (DOM) and methane ^14^. In oligotrophic high-As aquifers, DOM is a critical carbon and energy source, originating from (i) plant-derived material in aquifer sediments, (ii) modern surface-derived inputs (e.g., ponds, rivers, rice paddies, wastewater), and (iii) petroleum-derived hydrocarbons from thermally mature sediments ^15–19^. In addition to DOM, recent studies also implicate methane in driving As mobilization ^20,21^, particularly via anaerobic methane oxidation via reverse methanogenesis pathway coupled to Fe(III) reduction by members of the archaeal genus *Methanoperedens* ^20^. Microcosm incubations using natural wetland soils inferred that methane oxidation through reverse methanogenesis was linked to As(V) reduction, contributing significantly to arsenic release ^22^. In addition, *Methanosarcina acetivorans* has been shown to perform Fe(III)-dependent anaerobic methane oxidation via reverse methanogenesis, with acetate and CO_2_ as end products ^23^.

Although these findings point to a role for methane as an electron donor in arsenic-rich aquifers, the metabolic pathways and microbial interactions responsible for methane-driven Fe(III) and As(V) reduction remain poorly understood. This knowledge gap is especially relevant in methane-rich groundwater environments ^7,20,24,25^, where a complicated network of pathways may contribute to arsenic release. However, most experimental studies rely on complex sediment incubations, where diverse microbial communities can obscure the identification of key methanotrophic taxa and their metabolic roles.

To overcome these limitations, in-situ microbial capture techniques, such as “bio-traps” represent a promising new approach ^26^. These methods selectively enrich metal-reducing microbial communities in reducing aquifers ^27^ and, when combined with laboratory incubations and metagenomic analysis, offer a powerful approach to streamline investigations into Fe(III) and As(V) reduction processes and help uncover the mechanisms behind electron donor-Fe(III)/As(V) interactions.

In this study, we utilized Fe(III)-coated pumice stones as “microbial baits” to capture planktonic microbial communities from high-arsenic groundwater with elevated methane concentrations in the Kendal Province, Cambodia ^27–30^. After a 270-day field incubation, captured microbial communities were transferred to carefully-controlled anaerobic microcosms using methane or volatile fatty acids (VFAs) as electron donors. Genome-resolved metagenomics and stoichiometric analyses were used to assess microbial community function and electron flow. Our findings reveal that methane oxidation, primarily via reverse methanogenesis, led to acetate production, which served as a key intermediate driving Fe(III) and As(V) reduction. This work provides new insight into methane-linked biogeochemical cycling in arsenic-rich aquifers and its implications for arsenic mobilization and subsurface carbon cycling.

## Materials and Methods

### Collection and transport of pumice stones

Fe(III)-coated pumice stones were packed into acrylic sample holders with 2-mm perforations for water flow, and multiple holders were installed at the same depth in several boreholes at each of two hydrogeologically contrasting sites in Kendal Province, Cambodia: a clay-dominated site with average total dissolved As of 33 μg/L (ranged from 9.8 μg/L to 59 μg/L) and a sand-dominated site with average total dissolved As of 61 μg/L (ranged from 55 μg/L to 66 μg/L) (Fig. 1a). The groundwater geochemistry in these boreholes was published elsewhere ^31,32^. The detailed preparation of pumice stones, following an established method ^27^, and the in-situ installation are described in the Materials and Methods section of the Supporting Information. The stones (10 g) were collected from three boreholes from each site after nine months of incubation, and were stored in sterile 50 mL tubes (one tube from each well), with the tube topped up with filtered (through a 0.2 μm filter) groundwater collected from the same well. The tubes were stored on ice, transported to Manchester on ice and stored at 4°C prior to use. Pumice stones collected from all the boreholes at the same site (10 g per well) were combined before being used as an inoculum for the microcosm bottles as detailed below.

**Figure 1.**
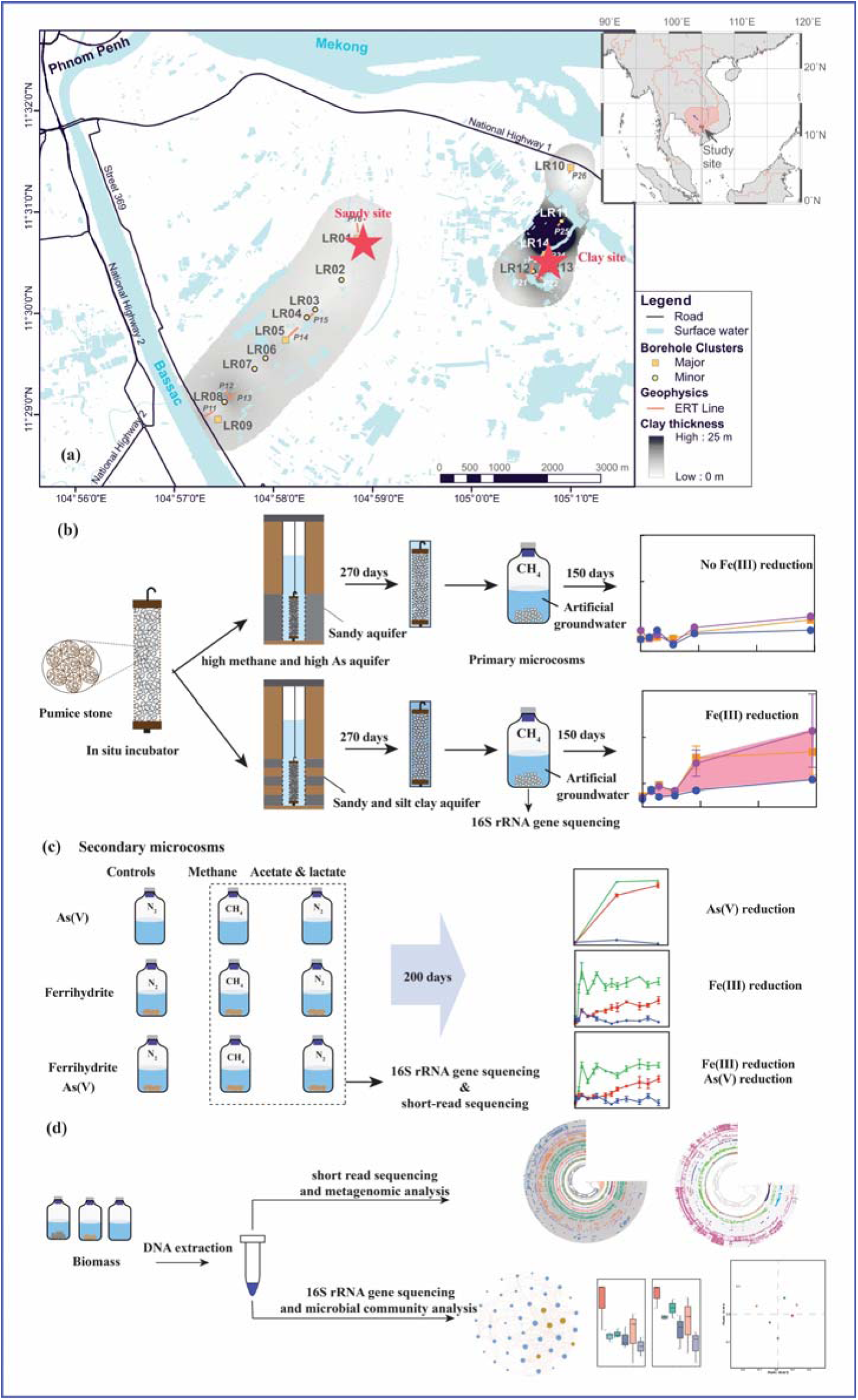
(a) Location of the clay-dominated and sand-dominated sites (red stars) in Kandal Province, Cambodia. LR marks transects of previous hydrogeochemical and electrical resistivity surveys in the area (from 2014 to 2015) ^30,80^. Adapted from Richards et al. (2019) ^39^; (b) the *in-situ* pumice stone installation and primary microcosms of pumice stone captured microbial communities; (c) primary microcosms of pumice stone captured microbial communities; and (d) the microbial community and metabolic potential identification.

### Primary microcosms preparation and sampling

Synthetic groundwater (1 L) was prepared for microcosm incubations as described previously ^33,34^, and was supplemented with 10 mmol/L synthetic ferrihydrite ^35^, or 10 mmol/L synthetic ferrihydrite and 0.1 mM sodium arsenate (Sigma-Aldrich, St. Louis, MO) as electron acceptors. The solutions were made anaerobic by flushing with an N_2_:CO_2_ gas mix (80:20 (v/v)) for 2 hours while monitoring the pH (the pH was adjusted to 7.0 by changing the flow of CO_2_). Inside the anaerobic chamber, each stock was split into 100 mL serum bottles containing 30 mL solution each and were sealed with butyl rubber stoppers before being sterilized via autoclaving. Triplicate microcosms were established for three treatments for each of the 2 sets: (i) no electron donor control, where the headspace was flushed with N_2_ for two minutes, (ii) CH_4_ as electron donor, where the headspace was flushed with CH_4_ for two minutes (the headspace of these incubations was replenished with CH_4_ after collecting the samples at the 98-day timepoint); (iii) VFAs as electron donors, where a solution of 5 mM sodium acetate and 5 mM sodium lactate was added as an electron donor and the headspace was flushed with N_2_ for two minutes (Fig. 1(a) and 1(b)). All glassware was prewashed with 5 % (v/v) nitric acid (HNO_3_, Sigma-Aldrich, St. Louis, MO) and milli-Q water before use. The bottles were inoculated with 1 g each of pumice stones that had been emplaced for nine months in boreholes in the Kendal Province, Cambodia. Note that two sets of laboratory incubations were prepared, one inoculated with pumice stones that had been deployed in the clay-dominated boreholes, and the other deployed in the sand-dominated boreholes. All microcosm bottles were incubated at 30 ^°^C for 150 days, and 1 mL samples were taken from the bottles using sterile syringes under anaerobic conditions for measuring the pH, and the concentration of Fe(II) in the microcosm slurries using the ferrozine assay ^36^. Experimental end-point samples were collected using the same procedure and were frozen at -20°C for DNA extraction and microbial community identification by 16S rRNA gene sequencing.

### Secondary microcosms preparation and sampling

The test microcosm bottles with methane as the electron donor that showed Fe(III) reduction were used to inoculate (10% v/v inoculum) secondary microcosms. The secondary microcosms were prepared following the same procedure as the primary microcosms, but an additional condition was also tested where 0.1 mM As(V) was added to the solution as the sole electron acceptor in addition to the two conditions that were tested in the primary microcosms (10 mmol/L ferrihydrite, and 10 mmol/L ferrihydrite and 0.1 mM As(V)) (Fig. 1(c)). The culture bottles were incubated at 30°C for 200 days, and 1 mL samples were taken from the bottles using sterile syringes under anaerobic conditions for measuring the pH, dissolved Fe(II) using the ferrozine assay ^36^, and dissolved As(III) and As(V) using high-performance liquid chromatography-inductively coupled plasma mass spectrometry (HPLC-ICP-MS) ^37^. Experimental end-point samples were collected using the same procedure and were frozen at -20°C for DNA extraction, microbial community identification by 16S rRNA gene sequencing, and metabolic potential by short-read metagenomic sequencing.

### Analytical measurements

The concentrations of volatile fatty acids (VFAs) were determined by ion exchange chromatography as described previously ^38^. The dissolved organic carbon (DOC) content in the liquid phase of the microcosm experiments was measured using the high-temperature catalytic oxidation method as described previously ^39^. The ratios of the two arsenic species were determined using HPLC-ICP-MS as described previously ^28,39^.

### Microbial community and metagenomic analysis

DNA was extracted from the frozen (at -20°C) samples using the ZymoBIOMICS DNA Miniprep Kit (Zymo Research, California, USA) following the manufacturer’s instructions. The extracted DNA was quantified using the Qubit dsDNA HS Assay Kit and a Qubit 3.0 Fluorometer (Invitrogen, Paisley, UK). The V4 hypervariable region of the 16S rRNA gene was amplified by polymerase chain reaction (PCR), prior to sequencing on an Illumina MiSeq platform (Illumina, USA) as described previously ^40^, along with appropriate DNA extraction and PCR controls. The demultiplexed fastq files were analysed using QIIME 2 version 2020.11^41^ as described previously ^40^, except that the Silva 138 99% reference database was used for taxonomic assignment. The produced amplicon sequence variant (ASV) tables, taxonomy tables, and phylogenetic trees were imported into R (version 4.1.0).

The DNA extractions were also used for building the metagenomic library and sequenced on an Illumina HiSeq 4000 platform (Illumina, USA) as described previously ^40^. Raw reads were trimmed and quality-filtered using Fastp (v0.19.7) ^42^. High-quality reads from single samples were assembled into scaffolds using metaSPAdes version 3.15.4 (--k-list 21, 33, 55, 77) ^43^ and all-samples were co-assembled utilizing MEGAHIT v1.2.9 with default parameters ^44^. The assemblies of each sample and co-assemblies were binned using metaWRAP (metabat2 v2.12.1, MaxBin2 v.2.2.6, and concoct v0.4.0) ^45–48^. The contamination and completeness of metagenome-assembled genomes (MAGs) was estimated using CheckM (v.1.0.11) ^49^ and dereplicated at 99% average nucleotide identity using dRep v3.4.0 ^50^ (-comp 50 -con 10 options). Final good-quality MAGs (completeness > 50% and contamination ≤ 10%) were kept for downstream analysis, including taxonomic assignment using GTDB-Tk v2.1.1 (database release 214) ^51^, metabolic potential annotation (METABOLIC_v4.0) ^52^ with updated As cycling gene database ^53^. Trees of good-quality MAGs were visualized and annotated with Interactive Tree Of Life (iTOL) ^54^. Heatmap of metabolic pathways and the MW stores (metabolic weight score) were visualized using TBtools ^55^.

## Results and Discussions

### Primary microcosm cultures with long-term *in-situ* incubated pumice stones exhibited Fe(III) reduction capability

The Fe(III)-coated pumice stones were deployed for nine months in high-As groundwater hosting clay-dominated (Clay-PS) and sand-dominated (Sand-PS) aquifers in Kandal Province (Cambodia), to enrich for Fe(III)-reducing microbial communities for laboratory incubations. These colonized pumice stones were added to “primary” microcosms to probe the interactions between the captured Fe(III)-reducing bacteria, ferrihydrite (with and without sorbed As(V))) and the potential electron donors (methane or lactate/acetate). Ferrihydrite supplemented microcosms inoculated with Clay-PS and Sand-PS colonized pumice baits and incubated without added electron donor, showed limited Fe(III) reduction (Fig. 2 and table S1 in Supporting Information), indicating little carryover of electron donors from the environment into the microcosm incubations. Microcosms with acetate and lactate as added electron donors and inoculated with Clay-PS and Sand-PS baits showed Fe (III) reduction to Fe(II) after 7 days of incubation (Fig. 2(a) and 2(d)). Fe(III) reduction was also observed in the microcosms that were inoculated with Clay-PS baits, and supplemented with either ferrihydrite or ferrihydrite/As, with methane supplied as the electron donor, however this was not the case in parallel methane supplemented Sand-PS microcosm incubations. The reduction of Fe(III) was more pronounced in lactate/acetate-treated Clay-PS microcosms, where the Fe(II)_tot_ and R_Fe(III)red_ were over three times higher than those of methane-treated Clay-PS microcosms as the electron donor, respectively (t-test, *p* < 0.027) (Figs. 2(a-b) and 2(d-e); Table S1 in Supporting Information). Previous studies showed that dissolved organic carbon levels in groundwaters from the clay-dominated site are around two times higher than those from the sand-dominated site ^33^, and could be released from clay layers under hydraulic disturbance (such as over-pumping) ^56–60^ leading to differentiated microbial communities involved in Fe(III) and As(V) reduction ^18,61,62^. The observed differences in Fe(III) reduction dynamics between the ferrihydrite-supplemented Clay-PS and Sand-PS controls may reflect site-specific variations in the relative abundance or activity of indigenous Fe(III)- and As(V)-reducing microorganisms enriched by the pumice stone baits. The pH values increased in the early stage of incubation (before 14 days) but decreased afterwards, likely due to the production of CO_2_ or VFAs during lacate/acetate or methane utilization (discussed in the following section). Our initial findings extend previous observations by demonstrating that pumice stones can enrich Fe(III)-reducing communities across contrasting sediment types, with evidence for utilization of both methane and lactate/acetate as electron donors.

**Figure 2.**
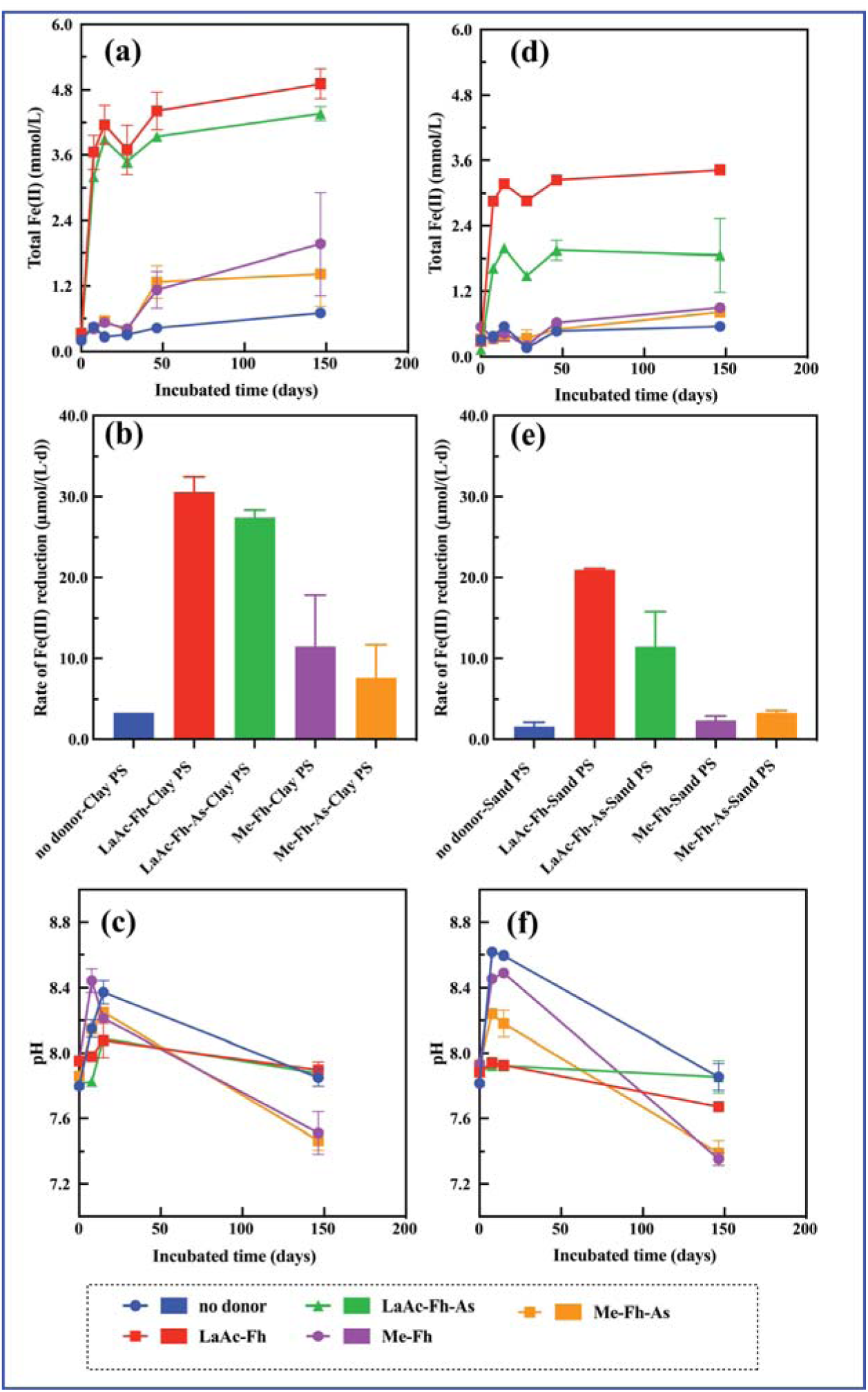
Fe(III) reduction kinetics ((a) and (d)), average rate of Fe(III) reduction ((b) and (e)), and changes of pH ((c) and (f)) of the microcosm samples that were prepared by supplementing artificial groundwater with electron acceptors (ferrihydrite and arsenate) and electron donors (LaAc, acetate and lactate; Me, methane). The samples that were inoculated with pumice stones previously incubated in the clay-dominated site (Clay-PS), and bottom panels represent the samples that were inoculated with pumice stones previously incubated in the sand-dominated site (Sand-PS) in Cambodia

After 150 days of microcosm incubation, microbial communities in the lactate/acetate-ferrihydrite Clay-PS microcosms were dominated by fermentative bacteria (*Anaerolineaceae*, 14.3%; *Acetothermiia*, 7.1%; *Latecibacteraceae*, 6.3%), Fe(III)-reducers (*Geobacteraceae*, 10%), and methanogenic archaea (*Methanobacteriaceae*, 5.9%; *Methanosaetaceae*, 5.8%; *Methanoregulaceae*, 4.9%) (Fig. S1). Community profiles were similar between lactate/acetate-ferrihydrite and lactate/acetate-ferrihydrite-As Clay-PS microcosms (Fig. S2). *Anaerolineaceae* have been associated with methanogenic archaea such as *Methanosaeta* in petroleum-impacted environments ^63^. At our study sites, petroleum-derived organic matter and elevated methane concentrations have been previously reported ^16^, potentially supporting syntrophic interactions between *Anaerolineaceae* and methanogenic archaea. Microbial communities in the methane-amended Clay-PS microcosms were dominated by fermentative *Anaerolineaceae* (21%) and members of the class *Thermodesulfovibrionia*, which increased from 4.9% in methane-ferrihydrite supplemented Clay-PS microcosms to 15.2% in methane-ferrihydrite-As Clay-PS, alongside Fe(III)-reducing *Geobacteraceae* (Fig. S1). The co-occurrence of *Thermodesulfovibrio* spp. and *Anaerolineaceae* spp. was previously detected in *n*-alkane-degrading methanogenic enrichment cultures, and were suggested to be generalist microbes participating in the metabolism of fermentation products ^64^. These observations also support the suggestion that petroleum-derived hydrocarbons from thermally mature sediments can support As mobilization from sediments into groundwaters in southeast Asia ^15–17,65^. Previous studies demonstrated the enrichment of Archaea belonging to the candidate genus *Methanoperedens* in long-term (220 days) microcosm cultures supplemented with methane, and inoculated with As-containing sediments from Vietnam ^20^. In our current study, methanogenic Archaea in general showed <1% abundance of the microbial community in the methane-amended Clay-PS samples (Fig. S1). Taken together, these results indicate potential syntrophic metabolic pathways driving methane oxidation and Fe(III) reduction.

### Secondary microcosm cultures acetate-coupled Fe(III) and As(V) reduction via methane oxidation

Secondary microcosm subcultures were prepared by adding a 10% (v/v) inoculum from the primary Clay-PS microcosm cultures to fresh artificial groundwater containing either ferrihydrite, ferrihydrite plus As(V), or soluble As(V) alone. In controls without added electron donors, no Fe(III) or As(V) reduction were detected and there was no production of VFAs (Figs. 3 and 4). In the As-supplemented secondary microcosm cultures, As(V) reduction was observed in the methane-As(V) enriched incubations (As(III)_tot_, 68 ± 3 μmol/L and R_As(V)red_, 0.35 ± 0.01 μmol/(L·d)) and complementary lactate/acetate-As(V) cultures (As(III)_tot_, 74 ± 1 μmol/L and R_As(V)red_, 0.38 ± 0.01 μmol/(L·d)). During As(V) reduction, lactate and acetate were consumed with propionate detected in the lactate/acetate supplemented As cultures (Fig. 4), suggesting fermentation processes alongside microbial As(V) reduction ^66^. Acetate accumulation was observed in the methane-As cultures (Ac_acu_, 96 ± 28 μmol/L; R_Acacu_, 0.49 ± 0.14 μmol/(L·d)) (Fig. 3), suggesting that methane oxidation have contributed to acetate production under these conditions.

**Figure 3.**
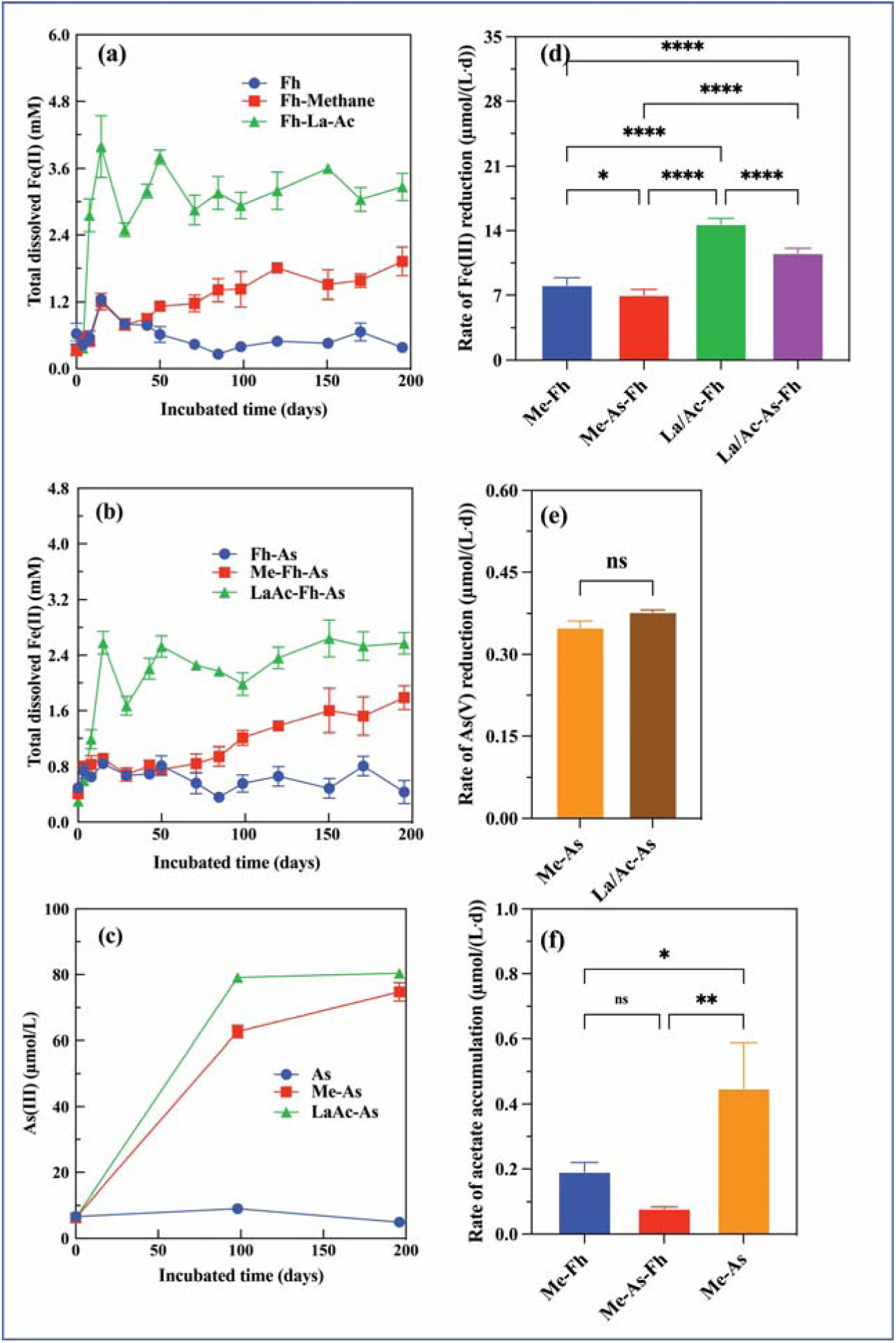
Fe(III) reduction kinetics and average rate of Fe(III) reduction in Fh treated enrichment cultures ((a) and (d)) and Fh-As treated enrichment cultures ((b) and (e)); As(V) reduction kinetics and average rate of As(V) reduction in As treated enrichment cultures ((c) and (f)). The differences of average rate of Fe(III) reduction or As(V) reduction were conducted using the one-way ANOVA test.

**Figure 4.**
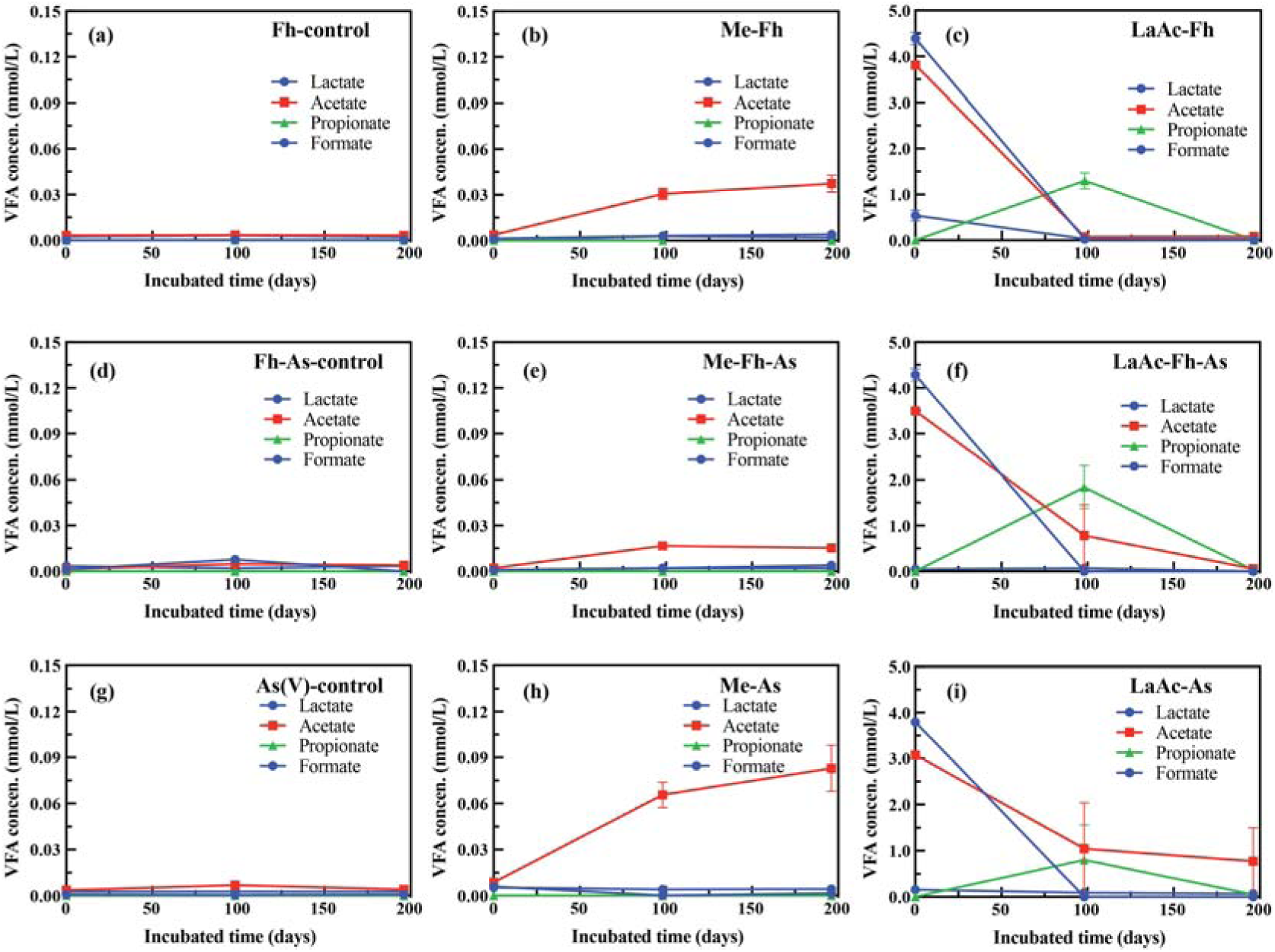
Versatile fatty acid analysis during the Fe(III) and/or As(V) reduction: in Fh-treated enrichment cultures ((a) to (c)), Fh-As-treated enrichment cultures((d) to (f)), and As-treated enrichment cultures((g) to (i))

In the ferrihydrite-supplemented secondary microcosm cultures, the added acetate and lactate were metabolized completely by day 98 of incubation in all treatments, leading to propionate production (ranging from 0.9 to 1.9 mmol/L), which was consumed by day 198 of the incubations (Fig. 4). Previous studies suggested that propionate can be produced by lactate fermentation and further converted to methane ^67^, while propionate degradation can be inhibited by acetate, resulting in propionate accumulation ^68^. Moreover, lactate/acetate additions resulted in higher rates of Fe(III) reduction and higher Fe(II)_tot_ levels than those noted in the methane-supplemented ferrihydrite cultures, indicating a higher efficiency of acetate and lactate utilization compared to methane oxidation for Fe(III) reduction. The presence of As(V) decreased the rate of Fe(III) reduction (and accumulation of Fe(II)_tot_) (Fig. 3 and Table S2 in Supporting Information). In the ferrihydrite–As(V) treated samples, total dissolved arsenic concentrations remained below the detection limit (0.13 nmol/L), likely due to arsenic sorption onto Fe-phases. In the presence of methane as the electron donor, the ferrihydrite-containing cultures showed a slow but steady reduction of Fe(III) to Fe(II) within 198 days of incubation (Fig. 3).This was accompanied by the production of acetate in the methane-ferrihydrite cultures (Ac_acu_,37 ± 6 μmol/L and R_Ac_, 0.19 ± 0.03 μmol/(L·d)) and methane-ferrihydrite-As cultures (Ac_acu_,15 ± 1 μmol/L and R_Ac_, 0.08 ± 0.01 μmol/(L·d)) (Fig. 4 and Table S2 in Supporting Information). Previous studies have suggested that acetate and CO_2_ could be the products of anaerobic methane oxidation driven by Fe(III)-dependent respiration ^23,69^. In our microcosms, the observed acetate accumulation, despite ongoing Fe(III) reduction, strongly suggests that acetate was produced as an intermediate during methane oxidation and utilized by Fe(III)-reducing microorganisms. These findings highlight a potential two-step process in which methane-derived acetate serves as a key intermediary linking methane oxidation to Fe(III) reduction under anoxic conditions.

After 198 days incubation, microbial communities clustered primarily according to the added electron donors followed by the electron acceptors (Figs. S2 and S3 in Supporting Information), which is expected given the specialized microbes that can use methane as a carbon and energy source under anaerobic conditions ^70^. Bacterial families known for fermentation (*Anaerolineaceae*, average relative abundance of 7.1%; and *Acidaminobacteraceae*, 1.0%) were noted in all secondary microcosm cultures, consistent with VFAs production, which was used during Fe(III)/As(V) reduction. Bacterial families associated with Fe(III) reduction (e.g. *Geobacteraceae*, average relative abundance of 17.5% and *Geothermobacteraceae*, 2.8%) were noted in all ferrihydrite-containing cultures, in line with the occurrence of Fe(III) reduction. It has been suggested that these specific groups of bacteria also contribute to As(V) reduction through respiration (if they express the *arrA* gene) or resistance mechanisms (for example by expressing the *arsC* gene) ^71^. Sulfate-reducing bacteria (*Syntrophobacteraceae* spp. and *Desulfurivibrionaceae* spp.) detected in these samples are also known to live under fermentation conditions in the absence of sulfate, which explains their detection in these incubations that have very low sulfate concentrations. Methanogenic archaeal families in these secondary cultures supplemented with methane were detected at >1% abundance (e.g. *Methanobacteriaceae* (6.6%), *Methanosaetaceae* (1.9%), and *Methanosarcinaceae* (1.4%)) in contrast to observations in corresponding primary cultures (Fig. S4 and Table S3 in Supporting Information). Methanogenic archaea may have been driving methanotrophic acetogenesis and the acetate produced could be used by other bacteria (e.g. Fe(III)-reducing and/or As(V)-reducing) and thus drive As release from As-containing sediments ^22,72–74^. This is an alternative or parallel mechanism to the previously suggested mechanism of direct coupling of anaerobic methane oxidation to Fe(III) reduction performed by members of the candidate genus *Methanoperedens* ^20^. Therefore, acetate production by methanotrophs via the reverse methanogenesis pathway may play an important role in Fe(III) and As(V) reduction within the enrichment cultures.

### Genome**-**resolved metagenomics revealed syntrophic metabolisms between methane oxidation and Fe(III) and As(V) reduction

To investigate the microbial pathways underlying methane-driven Fe(III) and As(V) reduction, we performed genome-resolved metagenomic analysis on the secondary microcosm cultures. A total of 153.5 Gb of high-quality sequence data (ranging from 21.9 to 32.0 Gb per library) were generated, yielding 264 medium-to-high quality metagenome-assembled genomes (MAGs) (completeness >50%, contamination ≤10%). These included 234 bacterial and 30 archaeal MAGs spanning 32 phyla, 54 classes, 72 orders, and 102 families (Fig. 4; Table S4 in Supporting Information). Among the bacterial MAGs, key groups included *Geobacteraceae* (9 MAGs), Desulfuromonadales spp. (5 MAGs), *Rhodocyclaceae* (11 MAGs), *Desulfarculaceae* (2 MAGs), *Melioribacteraceae* (11 MAGs), and *Anaerolineaceae* (20 MAGs), which together accounted for an average of 54 % of the microbial communities. Archaeal MAGs were dominated by *Methanotrichaceae* (11 MAGs), *Methanoregulaceae* (4 MAGs), *Methanosarcinaceae* (5 MAGs), *Methanobacteriaceae* (4 MAGs), *Methanocellaceae* (1 MAG), and UBA472 (3 MAGs), with an average abundance of 14 % (Fig. 5; Table S4 in Supporting Information).

**Figure 5.**
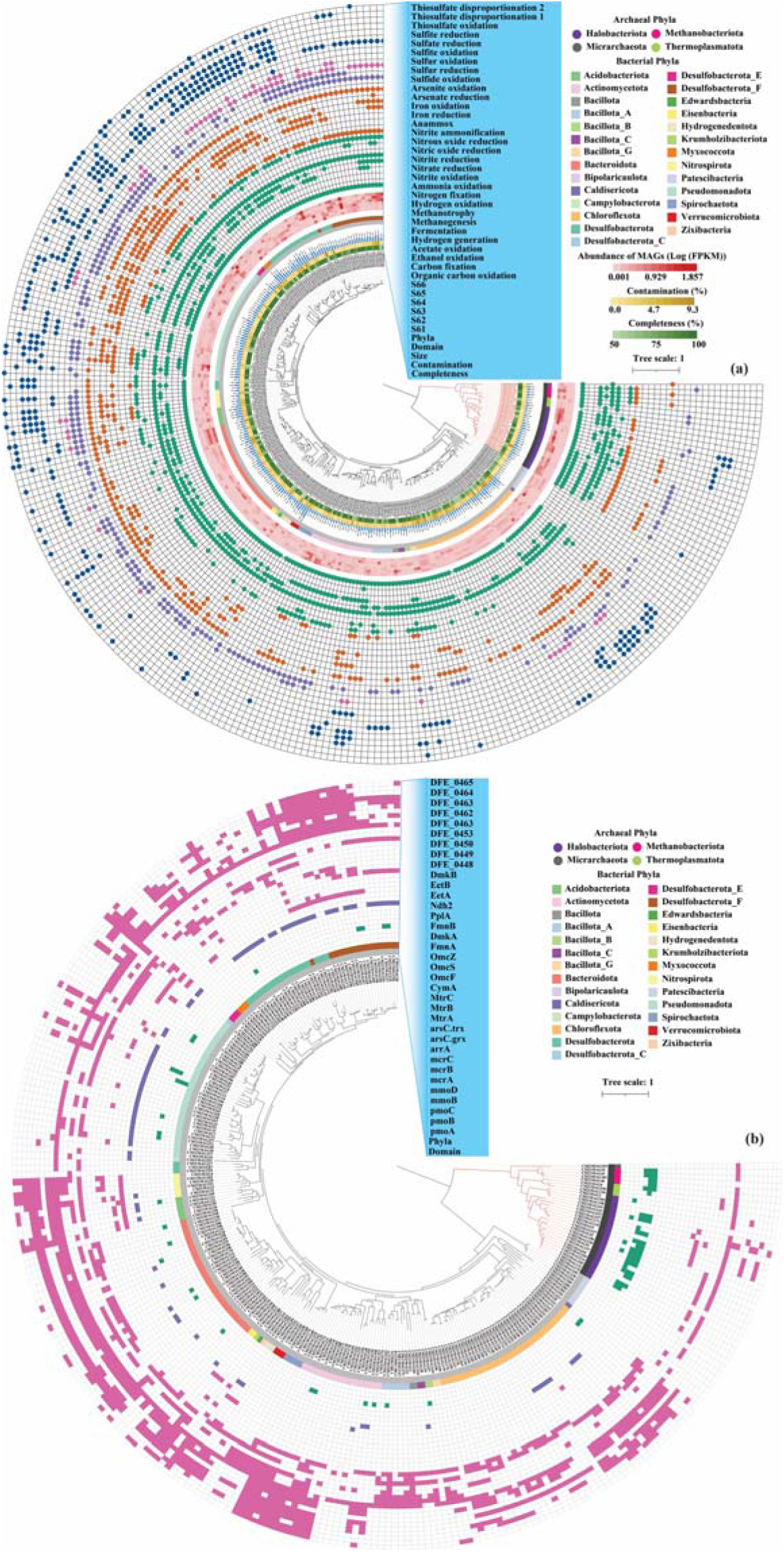
Genomic information, abundance, potential C-N-S-Fe-As metabolisms of the retrieved metagenomic assembled genomes (MAGs) (a) and functional genes related to fermentation to acetate, methanotroph, methanogenesis, Fe(III) reduction, and As(V) reduction (b).

Community-scale metabolic potential was assessed using metabolic weight (MW) scores derived from read-mapping and pathway coverage. Dominant functional pathways included fermentation (10 ± 1), acetate oxidation (6 ± 2), Fe(III) reduction (7 ± 1), As(V) reduction (4 ± 2), and sulfate reduction (3 ± 1) that were encoded in the microbial genomes (Table S5 in Supporting Information). Fermentation and acetate oxidation were the dominant carbon metabolic pathways for all secondary microcosm cultures, which could play a key role in supporting Fe(III) and As(V) reduction in the systems studied. Moreover, secondary microcosm cultures supplied with lactate/acetate or methane as electron donors, together with Fe(III) or Fe(III)-As(V) as electron acceptors, exhibited similar potential metabolic profiles. In contrast, cultures supplemented with only As(V) showed distinct metabolic patterns (Fig. S5; Tables S4 and S5, Supporting Information). Collectively, we conclude that the combination of electron acceptors (Fe(III) or As(V)) and electron donors (methane or lactate and acetate) were essential factors shaping the metabolisms of the microcosm cultures from the Clay PS-captured communities.

Genes or proteins associated with Fe(III) reduction were found widely in the retrieved genomes (257 MAGs), including those encoding *mtrABC* (38 MAGs, partial), *OmcS* (11 MAGs), *OmcF* (19 MAGs), *OmcZ* (7 MAGs), *FmnAB-dmkAB* (13 MAGs), *DFE_0448-0451* (19 MAGs), and *DFE_0461-0465* (24 MAGs) (Fig. 5b and Table S4 in Supporting Information), suggesting that diverse extracellular and intracellular pathways for Fe(III) reduction were represented in the secondary microcosm cultures. Moreover, the MW score for Fe(III) reduction did not significantly change across the Fe(III)-treated secondary microcosm cultures (6.33-7.42 across the cultures; t-test *p* values > 0.05; Table S5 in Supporting Information). Genes associated with As(V) respiratory (*arrA*) and detoxifying (*arsC*) reduction occurred in 86 bacterial MAGs (*arrA* encoded by 53 MAGs and *arsC* encoded by 33 MAGs), confirming the importance of bacterial As(V) reduction. MW scores of As(V) reduction were 1.75 in the methane-As cultures and 3.83 in the methane-ferrihydrite-As cultures. These were lower than those in lactate/acetate supplemented cultures (lactate/acetate-As(V), 4.17 and lactate/acetate-ferrihydrite-As(V) cultures, 6.47), inferring that either lactate/acetate or methane was used as electron donors for As(V) reduction in the secondary microcosm cultures.

Methane metabolism was assessed via MW scores and *mcr* gene distribution. In the lactate/acetate-amended cultures, MW scores for methanogenesis and methanotrophy were 2 ± 1 and 0.5 ± 0.8, respectively. In methane-only cultures, methanogenesis scores dropped significantly (0.06 ± 0.02; p = 0.026), and anaerobic methanotrophy was minimal (MW: 0.003 ± 0.001). This suggests that methane production was suppressed without organic substrates, and methane oxidation likely proceeded via alternative pathways. Notably, *mcr* genes linked to anaerobic methane oxidation via reverse methanogenesis were detected in three archaeal MAGs (CM21MAG237, CM21MAG172, and CM21MAG5), all affiliated with Methanosarcina. These MAGs were present in all treatments but showed the highest rpkm in the lactate/acetate-As(V) culture (38.91), moderate values in lactate/acetate-Fe(III) (1.16) and Fe(III)-As(V) cultures (0.89), and very low levels in methane-only treatments (0.041–0.12). This indicates that reverse methanogenesis occurred across conditions, as previously hypothesized^23^, but was most heavily represented in lactate/acetate-amended systems.

The genes encoding for enzymes for Fe(III) reduction, including *FmnAB-dmkAB*, *DFE_0448-0451*, and *DFE_0461-0465*, were found in two MAGs from the *Methanosarcina* genus (CM21MAG5 and CM21MAG172). Positive Spearman correlations were observed between the abundance of MAGs encoding *mcr* genes associated with methanogenesis and those encoding genes for Fe(III) and As(V) reduction (Fig. 6), further supporting a potential co-expression of these metabolic processes.

**Figure 6.**
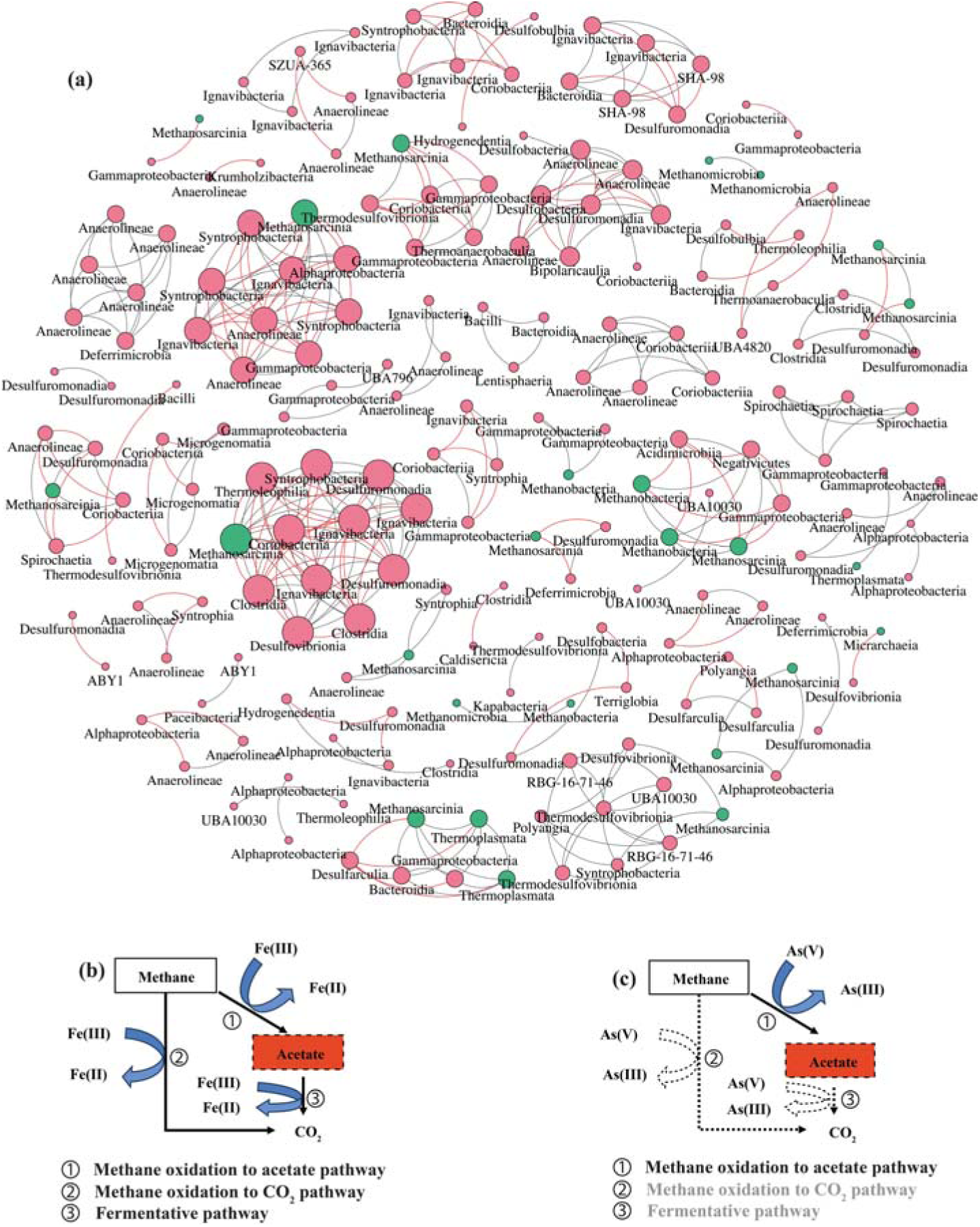
Spearman correlation network analysis of the retrieved metagenomic assembled genomes (MAGs) (a), pathways of methane oxidation in methane-treated ferrihydrite containing enrichment cultures (b), and pathways of methane oxidation in methane-treated enrichment cultures with As(V) as electron acceptors (b). Blue and red nodes in the network represent the archaeal and bacterial MAGs, respectively. Black edges indicate positive correlations while red edges indicate negative correlations. The solid-line black and dash-line grey arrows indicate the presence and absence of pathways of methane oxidation.

Interestingly, the propionate accumulation and subsequent utilization was observed in lactate/acetate treated series, while acetate continuously accumulated in the methane-treated series, with acetate accumulation higher in As(V)-methane treated systems. The *ack* and *pta* genes, identified in the genomes of CM21MAG172 and CM21MAG5, likely contribute to acetate production in both the methane-ferrihydrite and methane-As cultures. The presence of genes associated with reverse methanogenesis and Fe(III) reduction, alongside thermodynamic analyses, suggest the potential for multiple methane-oxidizing pathways, producing either acetate or CO_2_ as end products in these systems (Tables S4 and S6 in Supporting Information). However, considering that genes associated with As(V) respiratory (*arrA*) and detoxifying (*arsC*) reduction were not found in any of the archaeal MAGs, suggesting that methane-induced As(V) reduction is a syntrophic coupling of archaeal methane oxidation and bacterial arsenate reduction via acetate production and subsequent utilization. This model, via acetate production, provides a mechanistic explanation for the observed methane-driven As reduction. Taken together, our data support the hypothesis that methane drove As mobilization in the subsurface through direct reduction of As(V) and/or reduction of As(V)-bearing Fe(III) minerals, the latter with acetate acting as an intermediary in this process ^7,21,24^.

### Envrionmental Implications

Geogenic As contamination in reducing groundwaters is closely tied to interconnected biogeochemical processes, particularly the microbial reductive dissolution of As-bearing Fe(III) (oxyhydr)oxides. Although Fe(III) and As(V) reduction has been observed in diverse settings using various electron donors, including volatile fatty acids (VFAs) and methane, the key metabolic pathways driving these reductions, especially under methane-rich conditions, remain poorly understood. In the present study, we selected metal-reducing bacteria in groundwaters using Fe(III)-coated pumice baits, targeting boreholes that contained both dissolved organic matter and methane. Genome-resolved metagenomics, VFAs profiling and kinetic measurements of microbial As(V) and Fe(III) reduction in laboratory microcosms with the baits suggested that methane oxidation to acetate controlled microbial As(V) reduction by enrichment cultures, while multiple methane-oxidizing pathways with acetate as intermediate were involved in microbial Fe(III) reduction.

Our findings support a previously unrecognized pathway for methane-induced As mobilization in high-arsenic groundwater systems via the production of intermediate VFAs, which fuel downstream Fe(III) and As(V) reduction. This highlights the complexity of As-mobilizing processes in subsurface environments and expands current understanding of how microbial metabolisms are embedded within coupled biogeochemical networks. Although microbial communities with diverse functional traits that cooperate syntrophically are well-documented in oligotrophic subsurface aquatic ecosystems ^75–79^, the cross-feeding of methanotrophic acetate producers to Fe(III)-reducers and As(V)-reducers has not, to our knowledge, been previously implicated in As release in groundwater systems. This suggests that our findings may uncover a previously unexplored mechanism for As mobilization in subsurface environments. Notably, these pathways were only observed at a methane-bearing, clay-rich site, underscoring the importance of environmental context and the careful selection of microbial consortia. Further investigation is needed to elucidate the *in-situ* contribution of these methane-driven processes to As mobilization in Cambodian aquifers and other systems where such activities have been detected. By refining our understanding of both efficient “direct” VFA-driven pathways and the more complex, syntrophic methane-driven mechanisms of As release, we can better anticipate and manage the biogeochemical factors governing groundwater As enrichment.

## Supporting information

The Supporting Information is available free of charge with the electronic version of this paper.

## Notes

The authors declare no competing financial interest.

## Acknowledgements

This research has been supported by the Natural Environment Research Council (UK) (NE/P01304X/1 to Lloyd et al.). Dr. W.X. acknowledges support from the National Natural Science Foundation of China (W2511041), National Key Research and Development Program of China (grant No. 2021YFA0715902), and Fundamental Research Funds for the Central Universities (China) (grant no. 590223006). Chivuth Kong (formerly Royal University of Agriculture, now at Ministry of Agriculture, Forestry & Fisheries, Prey Veng Province, Cambodia) is thanked for field support. Dr. L.R. acknowledges support from a Dame Kathleen Ollerenshaw Fellowship and a UK Medical Research Council Future Leaders Fellowship (MR/Y016327/1). The local drilling team led by Hok Meas and local landowners are thanked for their contributions which enabled this work.

