## Supporting Information for "Methanotrophic acetogenesis drives a novel pathway of arsenic mobilization in reducing groundwaters"

**(total 7 pages including 5 figures and 6 tables)**

**Materials and Methods**

**The pumice stone preparation and installation**

The pumice stones of 4 – 6 mm in diameter were suspended in 37% HCl for 24 hours then washed repeatedly with deionised water until the runoff water pH became circumneutral. Acid-washed pumice stones were then mixed thoroughly with iron slurry at a ratio of 1:1 (w/v) and allowed to air-dry for 2 days, washed with deionised water to remove excess slurry, and then air dried for 3 days and stored dry for further use. The pumice stones were added to an acrylic sample holder with holes (2 mm diameter) to allow for water flow, and the holders were suspended in the wells at the top of the well screen (about 13 m depth from the top of the casing).

**Figure S1** Relative abundance of the 16S rRNA gene in primary microcosms that showed Fe(III) reduction to Fe(II), containing acetate and lactate or methane as electron donors, and ferrihydrite or ferrihydrite and As(V) as electron acceptors, after 150 days of incubation.

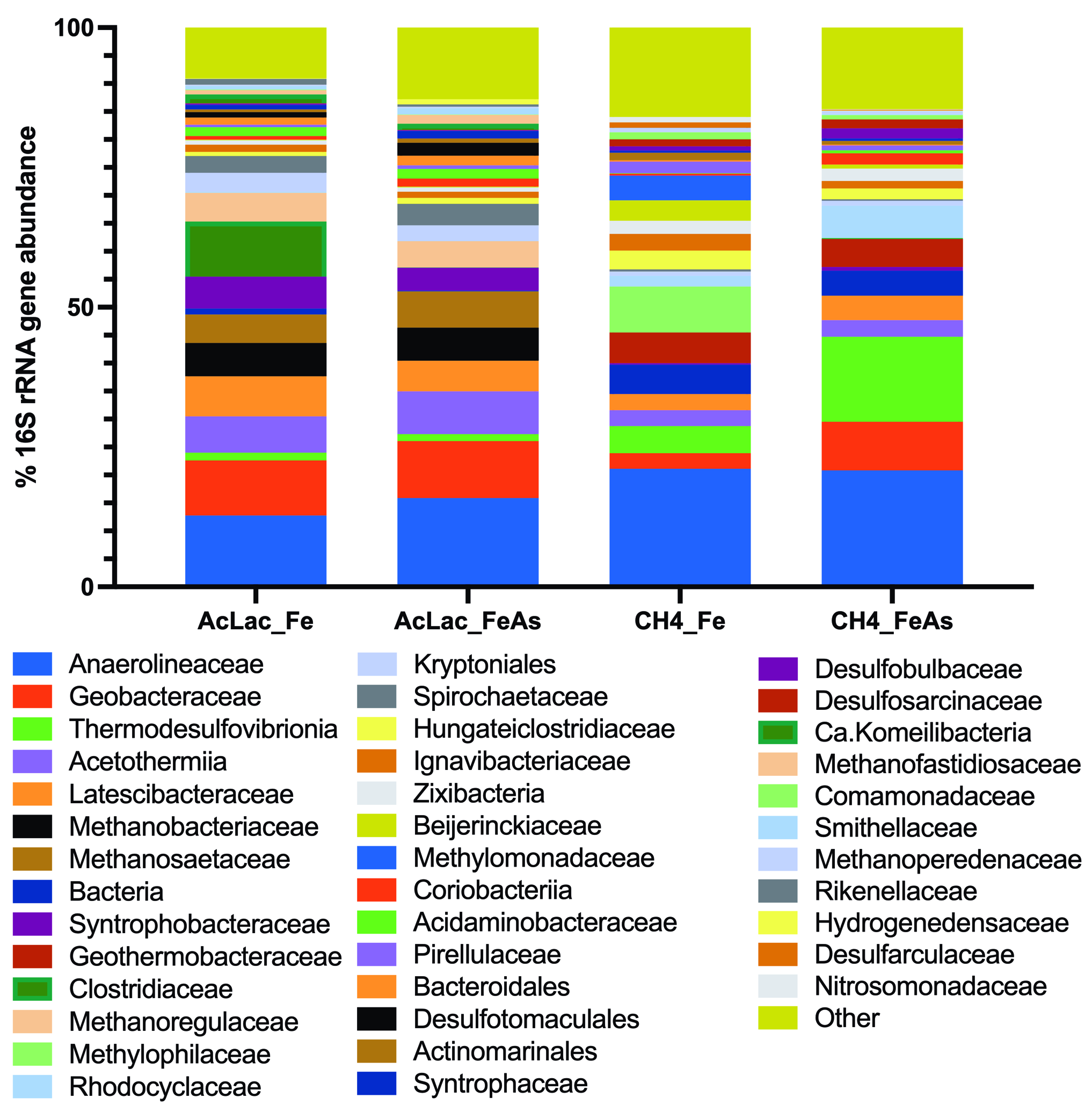

**Figure S2** Alpha diversity (Shannon and Simpson index) plot of the microbial communities in the secondary microcosms with LaAc or methane as electron donor. Within each group, triplicate samples are shown for each treatment (with As, Fh or Fh-As as electron acceptors).

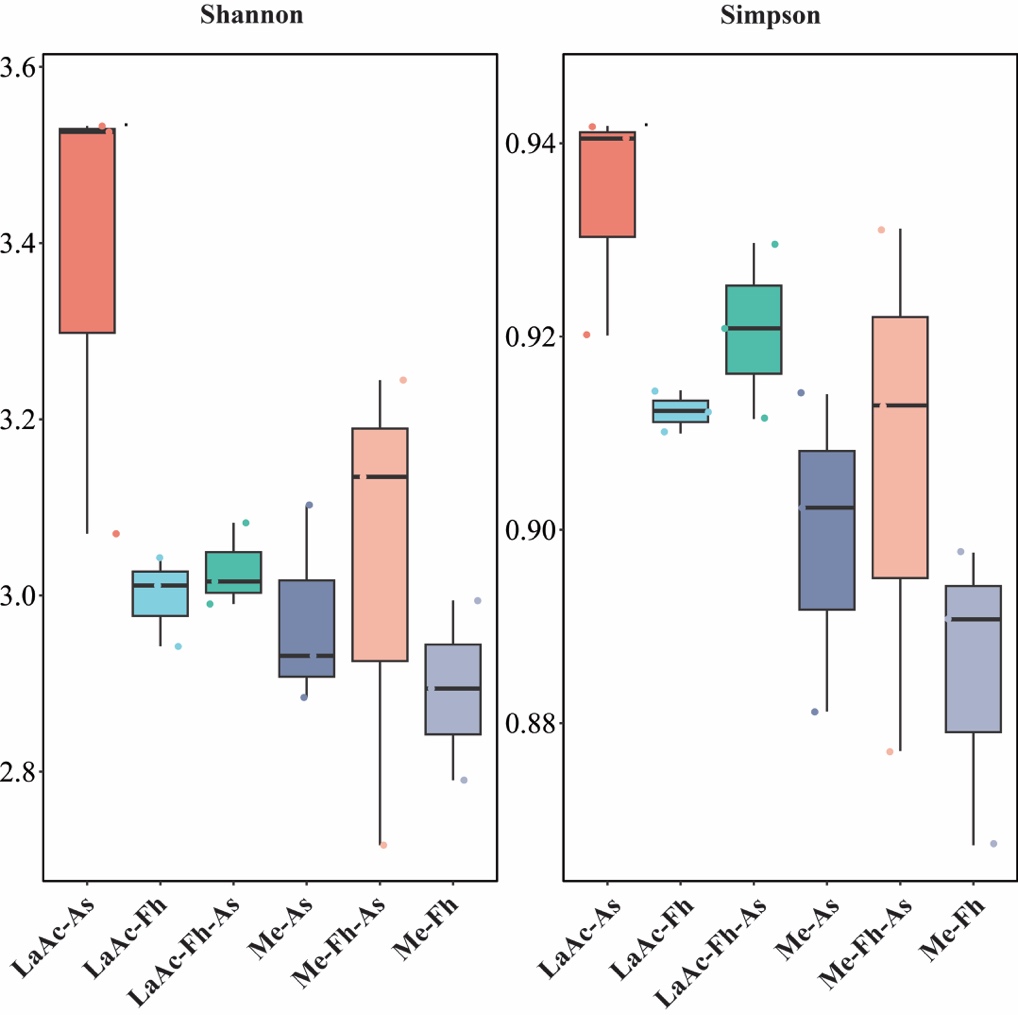

**Figure S3** Non-metric multi-dimensional scaling (NMDS) plot using Bray-Curtis dissimilarity matrix at the family level showing a stress of 0.116, and clustering of the samples from secondary microcosms based on the carbon source followed by the electron acceptor.

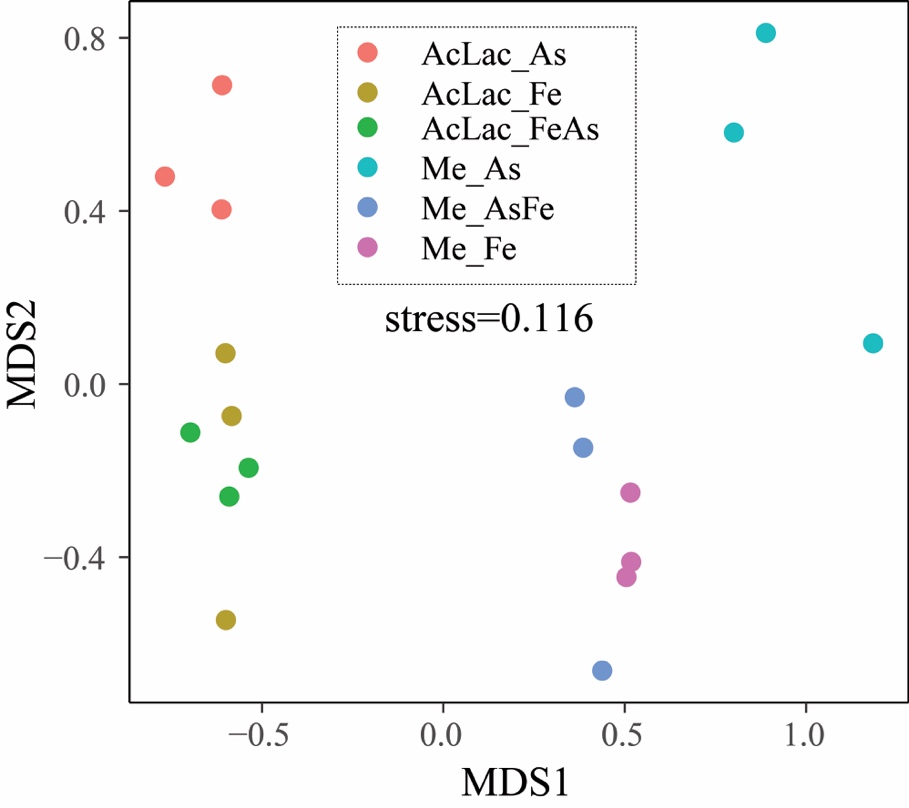

**Figure S4** Heatmap showing the microbial taxa that show >1% abundance of the 16S rRNA gene in at least one sample in the secondary microcosms.

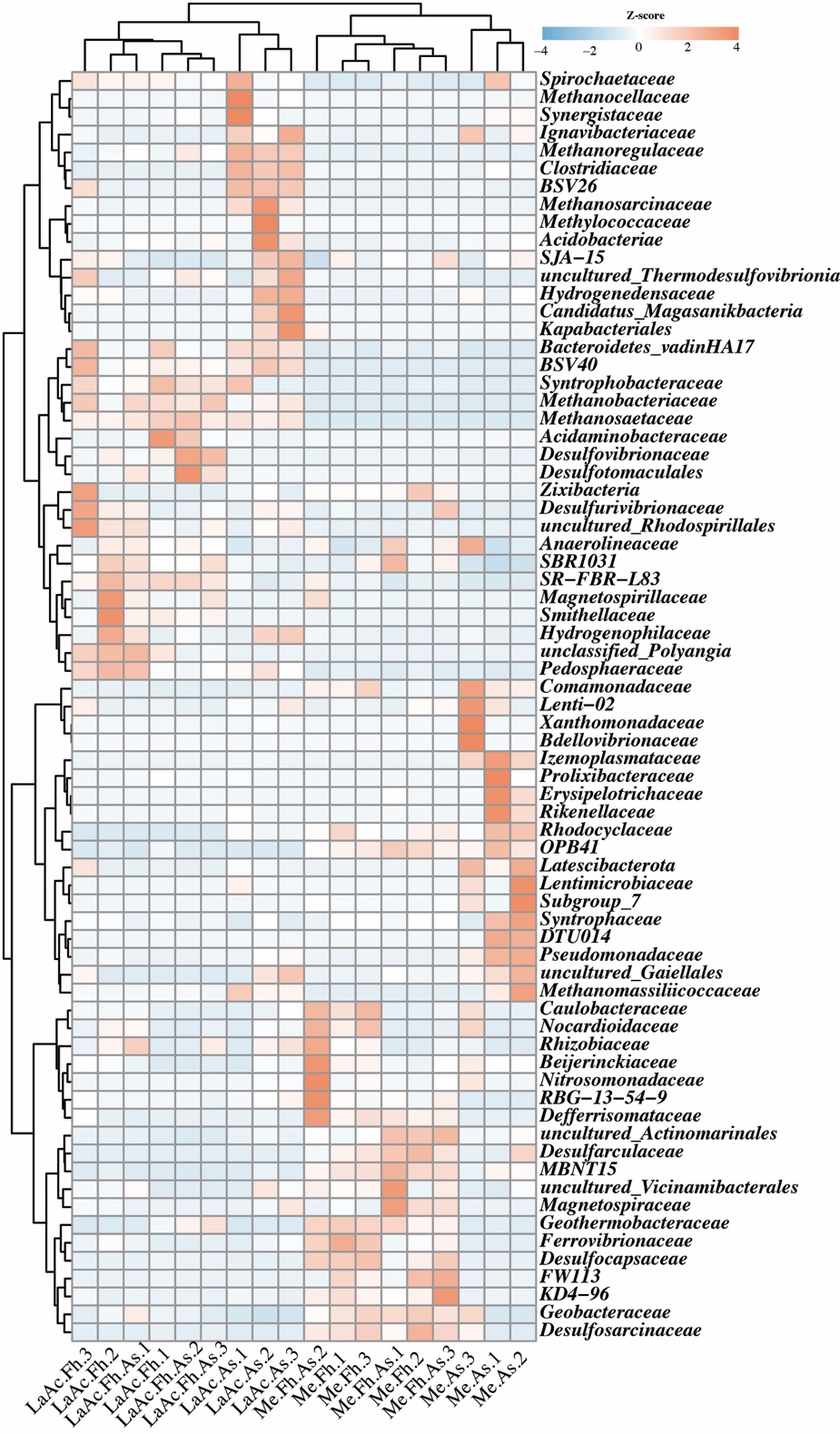

**Figure S5** PCoA plot using Bray-Curtis dissimilarity matrix based on MW scores of each metabolic pathway.

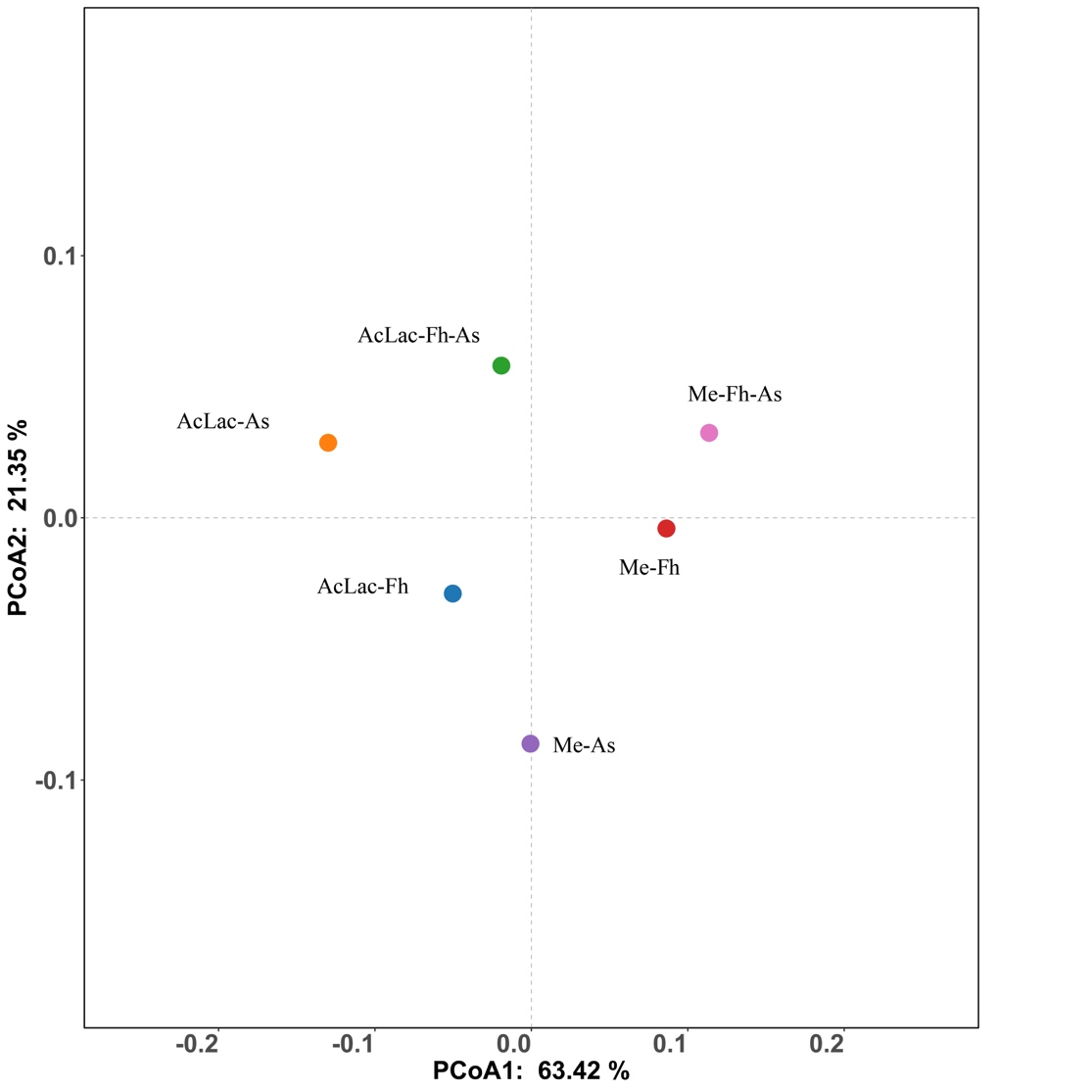

**Table S1** Total Fe(II) produced and rate of Fe(III) reduction of the primary microcosms with LaAc or methane as electron donor. Within each group, triplicate samples are shown for each treatment (with As, Fh or Fh-As as electron acceptors)

|  | Fe(II) production, mmol/L | Rate of Fe(III) reduction, µmol/(L･d) |
| --- | --- | --- |
| None electron donor Clay-PS | 0.50 ± 0.02 | 3.39 ± 0.45 |
| LaAc-Fh Clay-PS | 4.59 ± 0.28 | 31.29 ± 1.93 |
| LaAc-Fh-As Clay-PS | 4.12 ± 0.13 | 28.11 ± 0.89 |
| Me-Fh Clay-PS | 1.73 ± 0.95 | 11.76 ± 6.47 |
| Me-Fh-As Clay-PS | 1.14 ± 0.62 | 7.76 ± 4.22 |
| None electron donor Sand-PS | 0.24 ± 0.08 | 1.63 ± 0.55 |
| LaAc-Fh Sand-PS | 3.14 ± 0.02 | 21.44 ± 0.16 |
| LaAc-Fh-As Sand-PS | 1.72 ± 0.65 | 11.76 ± 4.43 |
| Me-Fh Sand-PS | 0.35 ± 0.08 | 2.41 ± 0.57 |
| Me-Fh-As Sand-PS | 0.49 ± 0.04 | 3.36 ± 0.30 |

**Table S2** Kinetic parameters of Fe(III) reduction, As(V) reduction, and acetate accumulation in the secondary microcosms with LaAc or methane as electron donor. Within each group, triplicate samples are shown for each treatment (with As, Fh or Fh-As as electron acceptors)

|  | Fe(II) production, mmol/L | Rate of Fe(III) reduction, µmol/(L･d) | As(III) production, µmol/L | Rate of As(V) reduction, µmol/(L･d) | Acetate accumulation, µmol/L | Rate of acetate accumulation, µmol/(L･d) |
| --- | --- | --- | --- | --- | --- | --- |
| As-control | *NA* | *NA* | *NA* | *NA* | *NA* | *NA* |
| Me-As | *NA* | *NA* | 68.45 ± 2.36 | 0.35 ± 0.01 | 87.41 ± 27.18 | 0.45 ± 0.14 |
| LaAc-As | *NA* | *NA* | 74.09 ± 0.68 | 0.38 ± 0.01 | *NA* | *NA* |
| Fh-control | *NA* | *NA* | *NA* | *NA* | *NA* | *NA* |
| Me-Fh | 1.59 ± 0.15 | 8.13 ± 0.79 | *NA* | *NA* | 37.33 ± 5.57 | 0.19 ± 0.03 |
| LaAc-Fh | 2.89 ± 0.12 | 14.80 ± 0.63 | *NA* | *NA* | *NA* | *NA* |
| Fh-As-control | *NA* | *NA* | *NA* | *NA* | *NA* | *NA* |
| Me-Fh-As | 1.59 ± 0.15 | 8.13 ± 0.79 | *NA* | *NA* | 15.36 ± 1.00 | 0.08 ± 0.01 |
| LaAc-Fh-As | 2.27 ± 0.10 | 11.63 ± 0.52 | *NA* | *NA* | *NA* | *NA* |

*NA* *indicated not available*

**Table S5** MW scores of each metabolic pathway calculated by using metabolic profiling and gene coverage from metagenomic read mapping at the community-scale level

| MW_score | LaAc-Fh | LaAc-As | LaAc-Fh-As | Me-Fh | Me-As | Me-Fh-As |
| --- | --- | --- | --- | --- | --- | --- |
| Organic_carbon_oxidation | 9.72 | 12.19 | 11.09 | 9.29 | 8.88 | 9.52 |
| Carbon_fixation | 2.60 | 2.17 | 2.22 | 0.74 | 2.51 | 0.35 |
| Ethanol_oxidation | 0.00 | 0.00 | 0.00 | 0.00 | 0.00 | 0.00 |
| Acetate_oxidation | 7.32 | 6.05 | 6.44 | 4.10 | 5.78 | 3.30 |
| Hydrogen_generation | 4.70 | 6.71 | 4.78 | 4.12 | 6.48 | 3.58 |
| Fermentation | 9.66 | 11.68 | 10.93 | 9.27 | 8.86 | 9.51 |
| Methanogenesis | 1.27 | 3.37 | 2.00 | 0.08 | 0.05 | 0.04 |
| Aerobic_methanotrophy | 0.00 | 0.22 | 0.00 | 0.00 | 0.06 | 0.00 |
| Anaerobic_methanotrophy | 0.04 | 1.39 | 0.03 | 0.00 | 0.00 | 0.00 |
| Hydrogen_oxidation | 5.85 | 5.35 | 6.06 | 7.91 | 6.40 | 8.47 |
| Nitrogen_fixation | 5.16 | 7.11 | 7.29 | 6.86 | 4.76 | 7.17 |
| Ammonia_oxidation | 0.00 | 0.00 | 0.00 | 0.00 | 0.00 | 0.00 |
| Nitrite_oxidation | 0.06 | 0.07 | 0.03 | 0.01 | 0.05 | 0.00 |
| Nitrate_reduction | 3.06 | 2.33 | 3.54 | 4.37 | 4.02 | 5.04 |
| Nitrite_reduction | 3.58 | 2.21 | 3.85 | 5.40 | 5.32 | 6.11 |
| Nitric_oxide_reduction | 3.44 | 2.60 | 1.56 | 3.61 | 3.41 | 3.02 |
| Nitrous_oxide_reduction | 2.27 | 3.20 | 1.62 | 1.19 | 1.58 | 0.87 |
| Nitrite_ammonification | 6.23 | 4.24 | 7.12 | 8.29 | 5.66 | 8.77 |
| Anammox | 0.00 | 0.00 | 0.00 | 0.00 | 0.00 | 0.00 |
| Iron_reduction | 6.33 | 4.97 | 7.42 | 7.02 | 6.19 | 7.30 |
| Iron_oxidation | 4.85 | 4.71 | 5.09 | 3.65 | 4.48 | 4.25 |
| Arsenate_reduction | 2.27 | 1.75 | 3.83 | 5.81 | 4.17 | 6.47 |
| Arsenite_oxidation | 0.73 | 0.64 | 0.68 | 0.22 | 0.11 | 0.23 |
| Sulfide_oxidation | 1.13 | 2.11 | 0.47 | 1.11 | 3.10 | 0.59 |
| Sulfur_reduction | 0.00 | 0.00 | 0.00 | 0.00 | 0.00 | 0.00 |
| Sulfur_oxidation | 4.64 | 4.62 | 5.48 | 6.89 | 6.15 | 7.06 |
| Sulfite_oxidation | 3.62 | 2.63 | 1.81 | 2.35 | 2.94 | 1.98 |
| Sulfate_reduction | 3.62 | 2.63 | 1.81 | 2.35 | 2.94 | 1.98 |
| Sulfite_reduction | 3.31 | 1.88 | 1.88 | 2.37 | 2.89 | 1.94 |
| Thiosulfate_oxidation | 0.18 | 0.88 | 0.16 | 0.24 | 2.34 | 0.10 |
| Thiosulfate_disproportionation_1 | 2.65 | 1.37 | 2.16 | 1.71 | 0.56 | 1.66 |
| Thiosulfate_disproportionation_2 | 1.72 | 0.91 | 0.64 | 1.03 | 0.29 | 0.67 |
